# Comparative transcriptomic analysis of maize ear heterosis during the inflorescence meristem differentiation stage

**DOI:** 10.1101/2021.09.28.462161

**Authors:** Xia Shi, Weihua li, Zhanyong Guo, Mingbo Wu, Xiangge Zhang, Liang Yuan, Xiaoqian Qiu, Ye Xing, Xiaojing Sun, Huiling Xie, Jihua Tang

**Affiliations:** National Key Laboratory of Wheat and Maize Crop Science, College of Agronomy, Henan Agricultural University, Zhengzhou 450002, China; Henan Institute of Crop Molecular Breeding, Henan Academy of Agricultural Sciences, Zhengzhou 450002, China

**Keywords:** Allele-specific expression, Heterosis, Inflorescence meristem, Maize (*Zea mays* L.), Nonadditive gene expression, Transcriptomics

## Abstract

Heterosis is widely used in many crops; however, its genetic mechanisms are only partly understood. Here, we sampled inflorescence meristem (IM) ears from the single-segment substitution maize (*Zea mays*) line lx9801^*hlEW2b*^, containing a heterotic locus *hlEW2b* associated with ear width, the receptor parent lx9801, the test parent Zheng58, and their corresponding hybrids. After transcriptomic analysis, 1638 genes were identified in at least one hybrid with nonadditively expressed patterns and different expression levels between the two hybrids. In particular, 2263 (12.89%) and 2352 (14.65%) genes showed allele-specific expression (ASE) in Zheng58 × lx9801 and Zheng58 × lx9801^*hlEW2b*^, respectively. A functional analysis showed that these genes were enriched in development-related processes and biosynthesis and catabolism processes, which are potentially associated with heterosis. Additionally, nonadditive expression and ASE may fine-tune the expression levels of crucial genes (such as *WUS* and *KNOX* that control IM development) controlling auxin metabolism and ear development to optimal states, and transcriptional variation may play important roles in maize ear heterosis. The results provide new information that increases our understanding of the relationship between transcriptional variation and heterosis formation during maize ear development, which may be helpful in clarifying the genetic and molecular mechanisms of heterosis.

## Introduction

Heterosis is an important method to improve crop yield and quality, and it plays a critical role in the breeding of several crops, including maize, rice, sorghum, and rape (Cheng et al., 2007; Kul’Pinova, 1966; Sernyk and Stefansson, 1983; Sokolov, 1970). However, the molecular mechanisms of heterosis are still not clear. At present, three classical hypotheses, dominance, overdominance, and epistasis, have been proposed to explain heterosis, and they have been debated for over 100 years (Lippman and Zamir, 2007). With advances in science and technology, a series of studies on heterosis using genomics, transcriptomics, proteomics, and epigenetics approaches have provided new insights into the molecular mechanisms of heterosis.

At the transcriptional level, gene expression variation causes changes in biological regulatory networks, which are important sources of phenotypic novelties and affect heterosis (Schadt et al., 2003). Comparing the differences in gene expression between parents and hybrids, multiple models of gene action, including additive, nonadditive, high- or low-parent dominance, and over- or underdominance, might be involved in the formation of heterosis (Swanson-Wagner et al., 2006). Several studies in maize (*Zea mays*) revealed that the additive effects are universal and have positive correlations with yield heterosis. In addition, the dominant and overdominant expression patterns, belonging to the nonadditive category, are also considered important factors of heterosis in hybrids (Rentao and Joachim, 2003; Song et al., 2007; Użarowska et al., 2007). The alterations in gene expression may cause changes in biological regulatory networks, which affect heterosis. By comparing the gene expression levels of hybrids and their parents at the maize ear developmental stage, Huang et al. (2006) found that most negative dominant genes are mainly involved in carbohydrate metabolism, lipid metabolism, energy metabolism, and protein degradation, whereas positive dominant genes are mainly involved in DNA replication and repair. In allotetraploid *Arabidopsis thaliana*, nonadditively expressed genes are significantly enriched in energy, metabolism, stress response, and plant hormone signal transduction (Jianlin et al., 2006).

In diploid hybrids, each gene contains two copies, one each from the male and female parents. Theoretically, the alleles from both parents should be equally expressed in the hybrid. However, the transcriptional activities of different alleles in hybrids vary greatly (Shi et al., 2012; Todesco et al., 2010). Allele-specific expression (ASE) refers to the preferential expression of a specific parental allele in its hybrids driven by regulatory factors from the parental genomes (Gaur et al., 2013). Hybridization produces an extremely large pool of allelic variants, which affect gene expression levels. The expression differences caused by ASE may lead to phenotypic diversity, depending on the gene functions (Guo et al., 2004). The ASE phenomenon has been documented in Arabidopsis, rice, maize, and barley (Guo et al., 2003; He et al., 2010; von Korff et al., 2009; Zhang and Borevitz, 2009). ASE patterns may have distinct implications in the genetic basis of heterosis, especially the dominance and overdominance hypotheses, because genetic variations frequently cause the differentially expression of genes, which may lead to phenotypic variations in the hybrids (Guo et al., 2006; Guo et al., 2004; Paschold et al., 2012; Springer and Stupar, 2007). Even though many genes in many species have been identified as exhibiting ASE at the whole-genome level, the potential relationship between ASE and heterosis remains unclear.

The important traits related to maize yield, such as kernel row number, kernel number per ear, ear width, and ear length, are all determined during inflorescence meristem (IM) development. The development of immature maize ears displays strong heterosis in ear architecture traits, which greatly affects grain yield (Jia et al., 2020). The size of the IM is significantly positively correlated with ear width and length, and its developmental process directly affects the final morphological characteristics of mature maize ears (Liu et al., 2015). The classic pathway of maintaining the IM amplification process is the *CLAVATA–WUSCHEL* (*CLV–WUS*) negative feedback loop. This pathway affects IM development by regulating the relationship between the proliferation of stem cells and the differentiation activities of tissues and organs (Somssich et al., 2016). *WUS* is a crucial regulator that determines stem cell formation and maintenance (Yadav et al., 2011). The *CLV3/WUS* negative feedback loop may affect the IM differentiation process (Brand et al., 2000). Additionally, analyses of mutants have established that the *BARREN STALK1* (*BA1)* gene encodes a *bHLH* transcription factor. Its mutant, *ba1*, cannot develop normal axillary meristem, and the proteins involved in auxin synthesis and transport do not function normally. It has been inferred that auxin deficiency may cause this mutants inability to complete the reproductive transition (Galli et al., 2015). *KNOTTED1-like homeobox* (*KNOX*), first isolated from maize, is involved in inflorescence development, as well as the maintenance and initiation of shoot apical meristem (SAM) and IM. *KNOX* regulates the dynamic conversion of SAM to IM by maintaining the balance of gibberellin and cytokinin, and its mutants may cause SAM malformation and the failure to complete the reproductive transformation process (Nathalie et al., 2012). Thus, hormones play critical roles in the regulation of immature maize ear development.

To eliminate the influence of genetic background and reduce environmental impact, single-segment substitution line populations have been developed to map heterotic loci. Chuanyuan et al. (2005) used single-segment substitution line test populations to research the heterosis performance of yield-related traits, and they discussed the advantages of single-segment substitution lines in heterosis research. Wang et al. (2012) detected 21 yield-related heterotic loci through the comparison of differences between a test cross population and background parents of 66 rice single-segment substitution lines. Wei et al. (2015) used the same experimental design to identify 21 heterotic loci related to maize plant architecture traits.

During the development of maize ear inflorescence, the IM stage is critical for ear development and heterosis. In the present study, we collected immature maize ears from single-segment substitution line lx9801^*hlEW2b*^ (containing the heterotic locus *hlEW2b* associated with ear width), receptor parent lx9801, test parent Zheng58, and their corresponding hybrids, Zheng58 × lx9801^*hlEW2b*^ and Zheng58 × lx9801, during the IM stage, and we developed a global gene expression profile using RNA sequencing (RNA-seq) technology to clarify the mechanisms of heterosis involved in transcriptome alterations. Our research provides new insights into the relationship between transcriptomic alterations and heterosis during maize ear development.

## Materials and methods

### Plant Material

The single-segment substitution line lx9801^*hlEW2b*^ containing an heterotic locus *hlEW2b* associated with ear width, receptor parent lx9801, and test parent Zheng58, as well as their corresponding test hybrids were grown in experimental fields in the summer of 2016 in Zhengzhou (Henan, China; E113° 65□, N34° 76□). In accordance with the leaf-age index, 2–4-mm immature maize ear samples were collected from each plant (only the first ear was collected per plant). A microscope was used to observe and determine the developmental period. Immature maize ears were manually collected at the IM differentiation stages. There were three biological replicates for each sample, and each biological replicate contains at least 30 immature maize ears. All the samples were immediately frozen in liquid nitrogen and stored at -80°C.

### RNA extraction and transcriptome sequencing

For the comparative transcriptome analysis, 15 IM samples (three replicates each for lx9801, lx9801^*hlEW2b*^, Zheng58, Zheng58 × lx9801, and Zheng58 × lx9801^*hlEW2b*^) were used to construct cDNAs and for RNA-seq. The total RNA of each sample was extracted with TRIzol® (Invitrogen, Carlsbad, CA, USA), and DNaseI was used to degrade the remaining DNA after extraction. The quality of the total RNA was determined using a Bioanalyzer 2100 system (Agilent Technologies, CA, USA). An Illumina TruSeq (Illumina, Santiago, CA, USA) RNA sample preparation kit was used to construct a library from each RNA sample. The prepared libraries were sequenced on an Illumina HiSeq2000 platform. Internal Perl scripts were used to remove linker sequences and low-quality sequences (sequences having lengths less than 120 bp) (Cole et al., 2012). After the original data were subjected to quality control measures, the paired-end clean reads were aligned to the maize B73 reference genome (Zea_mays.AGPv4.37) using Bowtie2 software (http://bowtie-bio.sourceforge.net/bowtie2/manual.shtml) with the following parameter: ‘-bowtie2-N 1’, which allows for only one base mismatch during the alignment process. Cufflinks (http://cole-trapnell-lab.github.io/cufflinks/) was used to assemble transcripts, and FPKM values were used to estimate gene expression levels.

### DEGs identification and gene expression pattern analysis

The DEGs between samples were estimated using the default parameters of Cuffdiff (Cufflinks software components). This software controlled the FDR of the *P*-value using the Benjamini and Hochberg method, and it set the differential expression threshold as FDR < 0.05. To gain overall insights into the gene expression patterns in the F_1_ hybrids, they were compared with MPVs. The threshold value was set as 0.05. Thus, FDR < 0.05 was considered as indicating a nonadditive expression pattern, and FDR > 0.05 was considered as indicating an additive expression pattern. As described by Ye et al. (2006), the GO (http://www.geneontology.org/) database was used for the functional analysis of the genes. The statistically enriched GO categories from the AgriGO2 website (http://systemsbiology.cau.edu.cn/agriGOv2/) were used, as was the single enrichment analysis (Singular enrichment analysis) tool to perform the statistically enriched GO categories analysis. The confidence criterion was FDR < 0.01.

### ASE identification

Transcriptome data were used to identify SNPs at the mRNA level between the parents. SAMtools (http://www.htslib.org/doc/1.2/samtools.html) was used to identify hybrid parental SNP information. The ‘-sort’ parameter of the SAMtools software was used to sort the output results of the Bowtie2 software and convert it into a ‘bam’ file. Parental ‘bam’ formatted files were used as input files and imported into the SAMtools software. The ‘mpileup’ and ‘bcftools’ parameters were used to identify SNP information between parents (Li et al., 2009a). In addition, the “-t DP” parameter was used to output the count results of the reference and nonreference genotypes of the SNP loci. Each parent was independently compared with the reference genotype to obtain the SNP count results between the parents.

RNAmodel is an R program based on hierarchical Bayesian model coding (http://cran.r-project.org/). It is used to identify ASE genes at the genome-wide level and to infer the results of allelic genotype counts for specific loci (Skelly et al., 2011). For the hybrids, the allele counts of each parent gene were imported into RNAmodel, and the alleles of both parents were quantitatively estimated to determine the expression levels of different alleles. Through a statistical analysis, the probability of ASE in each gene was determined, and ‘posterior probability’ was used to correct the results. The ASE identification criteria were as follows: 1) the specific expression level was greater than 0.8 or less than 0.4; 2) the posterior probability was greater than 0.8; and 3) there were at least two SNPs on one gene. Genes that met the above criteria were identified as ASE genes.

## Results

### Phenotypic analysis of near-isogenic lines and their corresponding hybrids

We previously identified the chromosome segment substitution line sub-CSSL_16_ carrying the ear-width heterotic locus *hlEW2b*, which contains a 1.98-Mb donor genomic segment from Chang7-2, and the phenotypic values of ear width and weight in Zheng58 × sub-CSSL_16_ are significantly higher than those of the control hybrid Zheng58 × lx9801 (*P* < 0.01). Simultaneously, the heterotic effects analysis revealed that the heterotic locus *hlEW2b* exhibits high overdominance (d/a ≥ 1, d = 4.19, a = 0.07) for heterosis (Shi et al., 2019). In the present study, the chromosome segment substitution line sub-CSSL_16_ containing the ear-width heterotic locus was redefined as lx9801^*hlEW2b*^. To evaluate the impact of the introgression locus for near-isogenic lines lx9801, lx9801^*hlEW2b*^ and their corresponding hybrids, Zheng58 × lx9801 and Zheng58 × lx9801^*hlEW2b*^, the phenotypic values of ear-related traits were investigated in multiple environments. In six environments, the phenotypic values of ear width and ear weight for Zheng58 × lx9801^*hlEW2b*^ were significantly higher than those of Zheng58 × lx9801, having average increases of 2.1 mm and 15.96 g, respectively, and their average over-standard heterotic values were 5.55% and 8.03%, respectively (Figure 1, Table 1). Additionally, there were no significant differences in the five ear traits between the inbred lines lx9801 and lx9801^*hlEW2b*^ in two environments (Supplementary Table S1).The results imply that there may be a heterotic locus associated with the Zheng58 allele in lx9801^*hlEW2b*^ that controlled the heterotic performances of ear width and weight in hybrid.

**Figure. 1.**
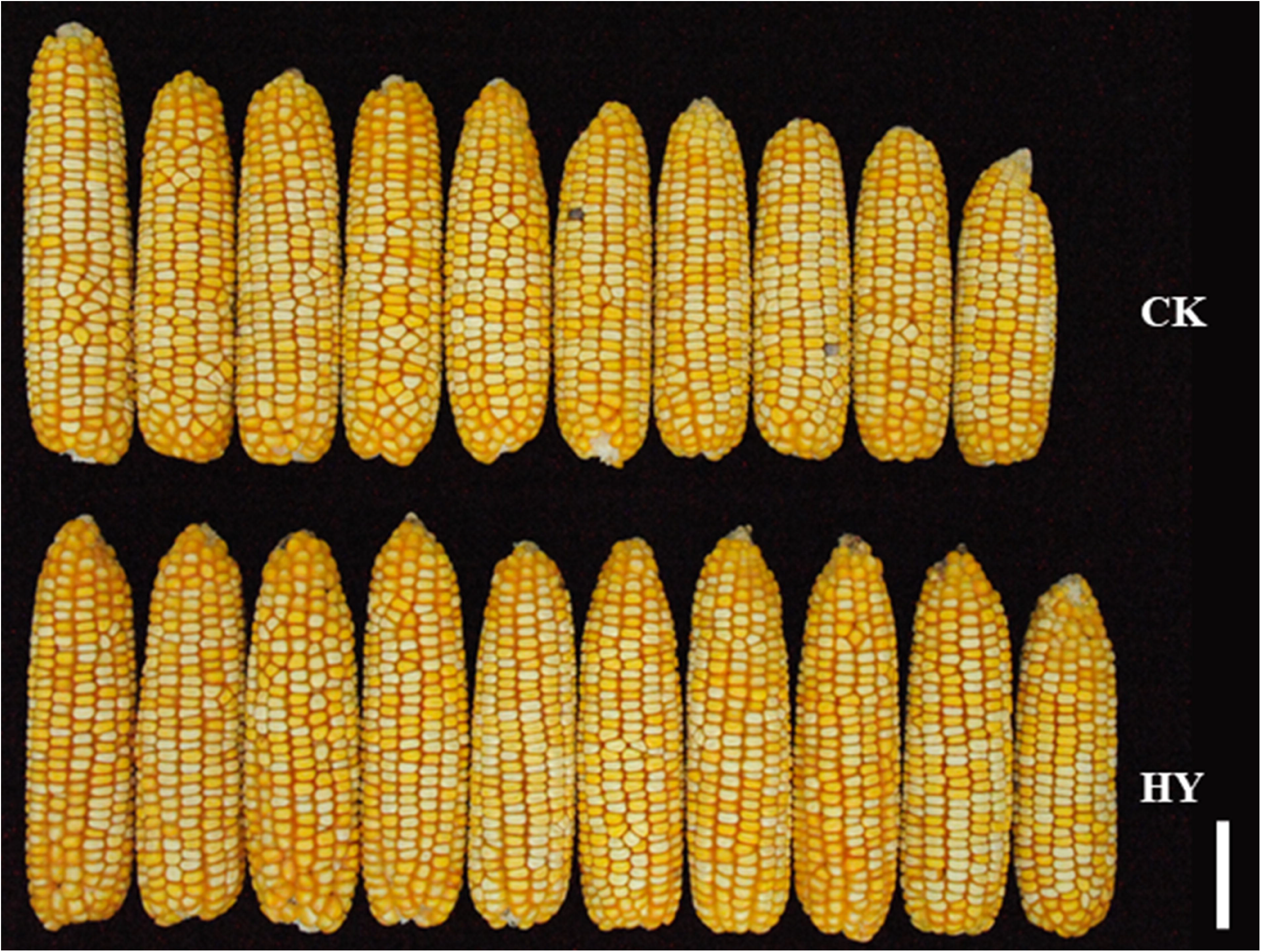
Heterosis performance of ear width trait between Zheng58×lx9801^*hlEW2b*^ and Zheng58×lx9801. The mature ear width traits of Zheng58 × lx9801^*hlEW2b*^ (HY) and Zheng58 × lx9801(CK), scale bar is 5cm.

**Table 1.**
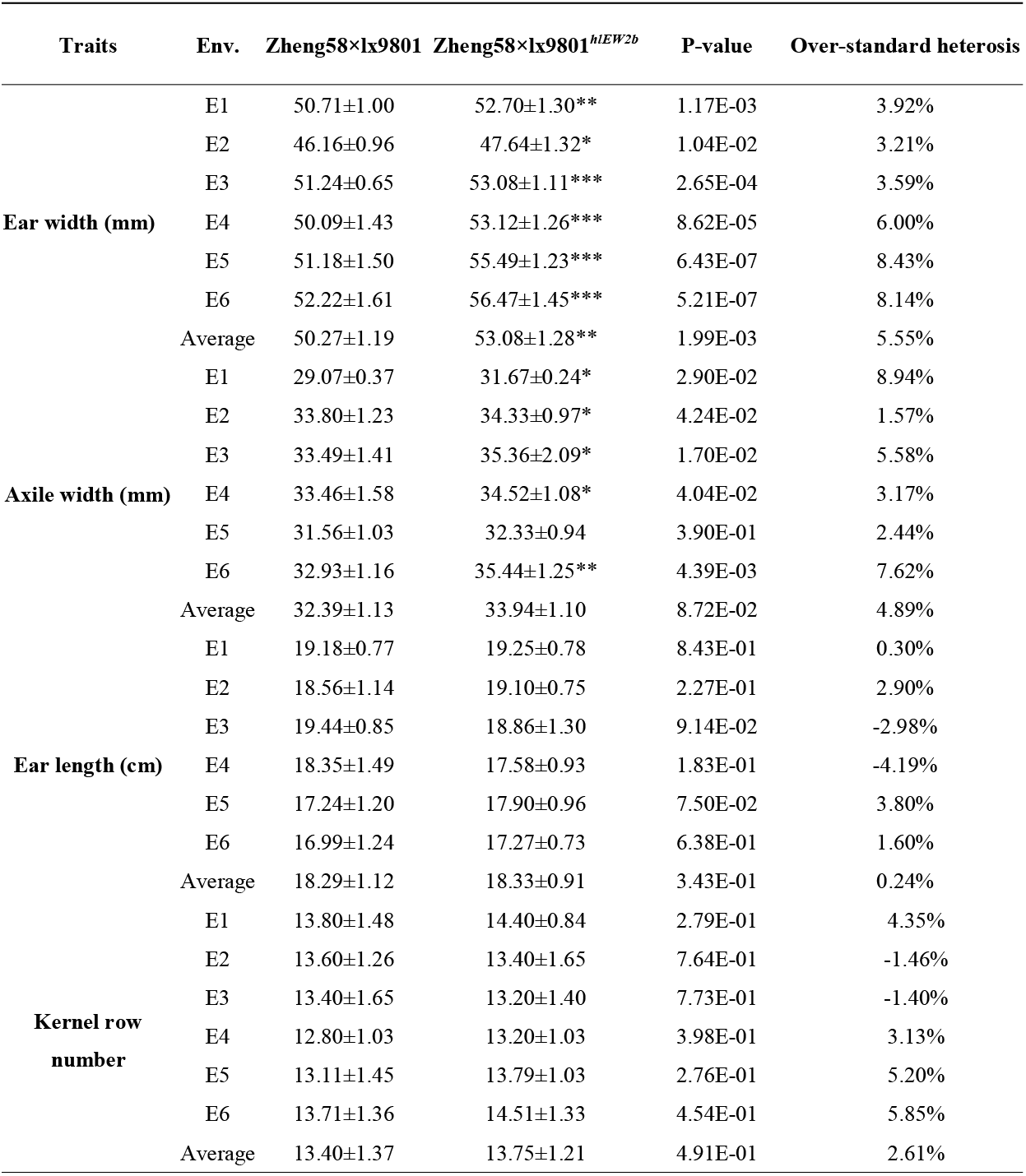
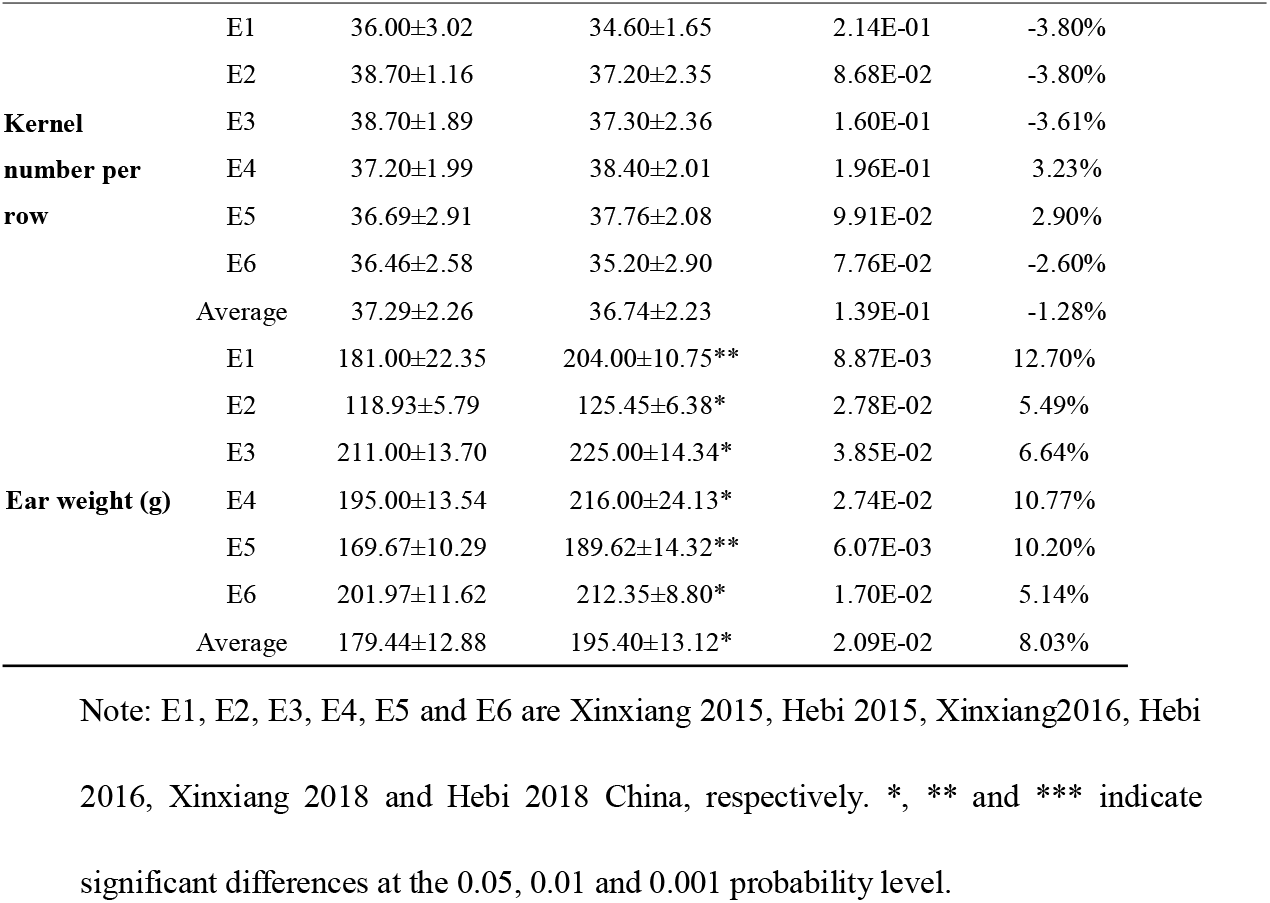
Comparison of ear traits between Zheng58 × lx9801 and Zheng58 × lx9801^*hlEW2b*^ in six environments.

| Traits | Env. | Zheng58×lx9801 | Zheng58×lx9801<sup>hlEW2b</sup> | P-value | Over-standard heterosis |
| --- | --- | --- | --- | --- | --- |
| Ear width (mm) | E1 | 50.71±1.00 | 52.70±1.30** | 1.17E-03 | 3.92% |
| | E2 | 46.16±0.96 | 47.64±1.32* | 1.04E-02 | 3.21% |
| | E3 | 51.24±0.65 | 53.08±1.11*** | 2.65E-04 | 3.59% |
| | E4 | 50.09±1.43 | 53.12±1.26*** | 8.62E-05 | 6.00% |
| | E5 | 51.18±1.50 | 55.49±1.23*** | 6.43E-07 | 8.43% |
| | E6 | 52.22±1.61 | 56.47±1.45*** | 5.21E-07 | 8.14% |
| | Average | 50.27±1.19 | 53.08±1.28** | 1.99E-03 | 5.55% |
| Axile width (mm) | E1 | 29.07±0.37 | 31.67±0.24* | 2.90E-02 | 8.94% |
| | E2 | 33.80±1.23 | 34.33±0.97* | 4.24E-02 | 1.57% |
| | E3 | 33.49±1.41 | 35.36±2.09* | 1.70E-02 | 5.58% |
| | E4 | 33.46±1.58 | 34.52±1.08* | 4.04E-02 | 3.17% |
| | E5 | 31.56±1.03 | 32.33±0.94 | 3.90E-01 | 2.44% |
| | E6 | 32.93±1.16 | 35.44±1.25** | 4.39E-03 | 7.62% |
| | Average | 32.39±1.13 | 33.94±1.10 | 8.72E-02 | 4.89% |
| Ear length (cm) | E1 | 19.18±0.77 | 19.25±0.78 | 8.43E-01 | 0.30% |
| | E2 | 18.56±1.14 | 19.10±0.75 | 2.27E-01 | 2.90% |
| | E3 | 19.44±0.85 | 18.86±1.30 | 9.14E-02 | -2.98% |
| | E4 | 18.35±1.49 | 17.58±0.93 | 1.83E-01 | -4.19% |
| | E5 | 17.24±1.20 | 17.90±0.96 | 7.50E-02 | 3.80% |
| | E6 | 16.99±1.24 | 17.27±0.73 | 6.38E-01 | 1.60% |
| | Average | 18.29±1.12 | 18.33±0.91 | 3.43E-01 | 0.24% |
| Kernel row number | E1 | 13.80±1.48 | 14.40±0.84 | 2.79E-01 | 4.35% |
| | E2 | 13.60±1.26 | 13.40±1.65 | 7.64E-01 | -1.46% |
| | E3 | 13.40±1.65 | 13.20±1.40 | 7.73E-01 | -1.40% |
| | E4 | 12.80±1.03 | 13.20±1.03 | 3.98E-01 | 3.13% |
| | E5 | 13.11±1.45 | 13.79±1.03 | 2.76E-01 | 5.20% |
| | E6 | 13.71±1.36 | 14.51±1.33 | 4.54E-01 | 5.85% |
| | Average | 13.40±1.37 | 13.75±1.21 | 4.91E-01 | 2.61% |

| Traits | Env. | Zheng58×lx9801 | Zheng58×lx9801<sup>hIEW2b</sup> | P-value | Over-standard heterosis |
| --- | --- | --- | --- | --- | --- |
| Kernel<br>number per<br>row | E1 | 36.00±3.02 | 34.60±1.65 | 2.14E-01 | -3.80% |
|  | E2 | 38.70±1.16 | 37.20±2.35 | 8.68E-02 | -3.80% |
|  | E3 | 38.70±1.89 | 37.30±2.36 | 1.60E-01 | -3.61% |
|  | E4 | 37.20±1.99 | 38.40±2.01 | 1.96E-01 | 3.23% |
|  | E5 | 36.69±2.91 | 37.76±2.08 | 9.91E-02 | 2.90% |
|  | E6 | 36.46±2.58 | 35.20±2.90 | 7.76E-02 | -2.60% |
|  | Average | 37.29±2.26 | 36.74±2.23 | 1.39E-01 | -1.28% |
| Ear weight (g) | E1 | 181.00±22.35 | 204.00±10.75** | 8.87E-03 | 12.70% |
|  | E2 | 118.93±5.79 | 125.45±6.38* | 2.78E-02 | 5.49% |
|  | E3 | 211.00±13.70 | 225.00±14.34* | 3.85E-02 | 6.64% |
|  | E4 | 195.00±13.54 | 216.00±24.13* | 2.74E-02 | 10.77% |
|  | E5 | 169.67±10.29 | 189.62±14.32** | 6.07E-03 | 10.20% |
|  | E6 | 201.97±11.62 | 212.35±8.80* | 1.70E-02 | 5.14% |
|  | Average | 179.44±12.88 | 195.40±13.12* | 2.09E-02 | 8.03% |

Note: E1, E2, E3, E4, E5 and E6 are Xinxiang 2015, Hebi 2015, Xinxiang2016, Hebi 2016, Xinxiang 2018 and Hebi 2018 China, respectively. \*, \*\* and \*\*\* indicate significant differences at the 0.05, 0.01 and 0.001 probability level.

To understand the developmental basis of ear width, the IM sizes of 2–4-mm immature ears were observed. The average diameter of ear IM in Zheng58 × lx9801^*hlEW2b*^ was 501.21 ± 19.98 μm, which was significantly larger (*P*-value = 3.36E-09) than that of Zheng58 × lx9801 (465.29 ± 17.04 μm) in the developing female inflorescence (Figure 2A, B). However, there were no significant differences in the lengths of IMs (Figure 2A, C), which was determined using the final lengths of mature ears between the hybrids Zheng58 × lx9801 and Zheng58 × lx9801^*hlEW2b*^ (*P*-value = 1.70E-01). Thus, the presence of the heterotic locus *hlEW2b* may significantly increase the widths of IMs in hybrids and may provide more space for ear development by increasing the sizes of the IMs, leading to the significantly greater ear width and weight of Zheng58 × lx9801^*hlEW2b*^ compared with the control hybrid Zheng58 × lx9801.

**Figure. 2.**
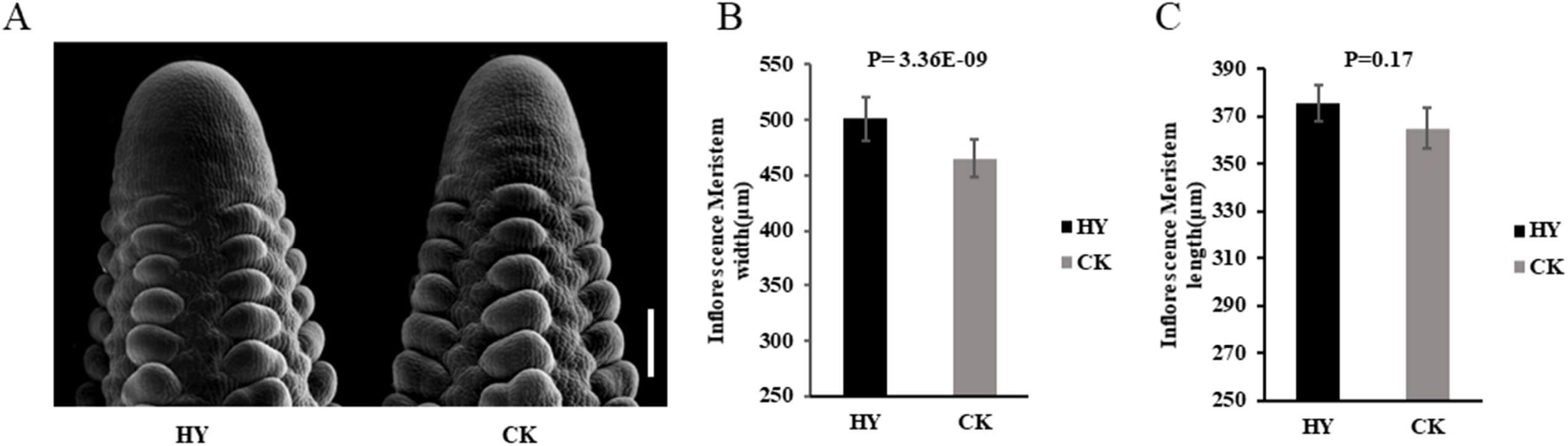
Inflorescence meristems size in hybrids. A: Scanning electron micrograph image of Zheng58×lx9801^*hlEW2b*^ (left) and Zheng58×lx9801 (right) 2-4mm immature ears, bar=200μm. B-C: Comparison of inflorescence meristems size between Zheng58×lx9801^*hlEW2b*^ and Zheng58×lx9801.

### Differential gene expression pattern analysis for heterosis

Based on the criteria that FPKM ≥ 1 in at least one genotype and false discovery rate (FDR) < 0.05, 2531 genes (59.8%) were differentially expressed only in Zheng 58 × lx9801^*hlEW2b*^ vs. Zheng 58 × lx9801, and 1303 significant differentially expressed genes (DEGs) were identified between lx9801^*hlEW2b*^ and lx9801. A Venn diagram revealed 400 genes (9.4%) were differentially expressed in both comparison groups. The comparison of DEGs in Zheng 58 × lx9801^*hlEW2b*^ vs. Zheng 58 × lx9801 was significant because the direct influence of the substitution fragment itself in the resulting hybrids was eliminated, thereby retaining the heterotic effects of the substitution fragment (Figure 3A). These DEGs may be important factors in the phenotypic differences between hybrids.

**Figure. 3.**
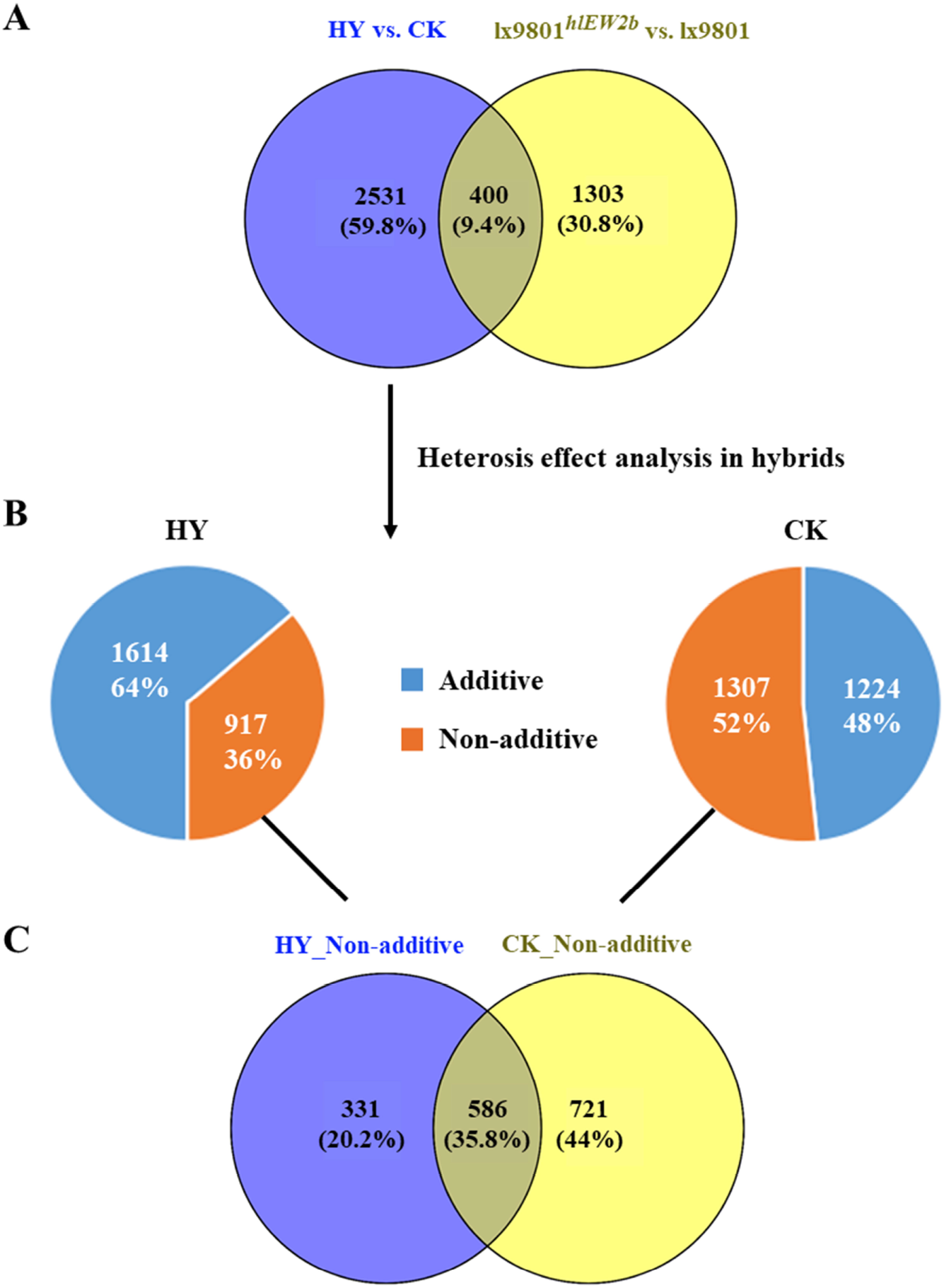
Expression pattern analysis of differentially expressed genes. A: Statistical analyses of differentially expressed genes among hybrids [Zheng58 × lx9801^*hlEW2b*^ (HY), Zheng58 × lx9801(CK)] and their corresponding inbred lines lx9801^*hlEW2b*^ and lx9801. B: Expression patterns analysis of specific differentially genes between hybrids. C: Venn diagram showing the number of non-additive expression genes selected from each hybrid.

To gain overall insights into the expression patterns of unique DEGs in Zheng 58 × lx9801^*hlEW2b*^ vs. Zheng 58 × lx9801, the additive and nonadditive patterns determined using pairwise comparisons between the hybrids and their corresponding mid-parent value (MPV) were further classified. A total of 1614 (64%) and 1224 (48%) genes were expressed additively (F_1_ vs. MPV, FDR > 0.05) in Zheng 58 × lx9801^*hlEW2b*^ and Zheng 58 × lx9801, respectively. In total, 917 and 1307 genes displayed nonadditive expression patterns (F_1_ vs. MPV, FDR < 0.05) in Zheng 58 × lx9801^*hlEW2b*^ and Zheng 58 × lx9801, respectively (Figure 3B). The presence of additively expressed genes in hybrids implied that there were additive effects on gene expression in hybrids, and nonadditively expressed genes may be the main contributors to heterosis.

### Functional characterization of nonadditively expressed genes

To ascertain the differences in heterotic performances between the two hybrids, we combined 917 and 1307 nonadditively expressed genes from Zheng58 × lx9801^*hlEW2b*^ and Zheng58 × lx9801 into one set of 1638 genes, in which each gene showed a nonadditive expression pattern in at least one hybrid combination and displayed different heterotic performances between Zheng58 × lx9801^*hlEW2b*^ and Zheng58 × lx9801 (Figure 3C). The nonadditively expressed genes in the union set were classified into the following three categories: 1) 586 genes (35.8%) showing nonadditive expression patterns in both Zheng 58 × lx9801^*hlEW2b*^ and Zheng 58 × lx9801 hybrids; 2) 331 genes showing a nonadditive expression pattern (20.2%) in Zheng 58 × lx9801^*hlEW2b*^ but not in Zheng 58 × lx9801; and 3) 721 genes showing a nonadditive expression pattern in Zheng 58 × lx9801 but not in Zheng 58 × lx9801^*hlEW2b*^ (Figure 3C). The three types of genes might be responsible for the different in heterotic performances between hybrids of Zheng 58 × lx9801^*hlEW2b*^ and Zheng 58 × lx9801.

A gene ontology (GO) enrichment analysis was conducted using the single enrichment analysis on the AgriGO website to determine the functional categories for nonadditively expressed genes from hybrids Zheng 58 × lx9801^*hlEW2b*^ and Zheng 58 × lx9801. These nonadditive DEGs were found to be enriched (FDR < 0.01) in the three hierarchically structured GO terms, molecular function (MF), biological process (BP), and cellular component (CC). The MF-related genes were enriched in six main GO terms, with the three most significant subcategories being RNA-directed DNA polymerase activity (GO:0003964, FDR = 1.10E-06), nuclease activity (GO:0004518; FDR = 6.60E-05), and nucleotidyltransferase activity (GO:0016779; FDR = 9.40E-05). Additionally, auxin binding-related genes were also significantly enriched (GO:0010011; FDR = 2.80E-03) (Figure 4A, Supplementary Table S3). In the BP category, among the top five GO items with the highest significance levels, two were related to growth and development, namely tissue development (GO:0009888; FDR = 3.70E-06) and plant organ development (GO:0099402; FDR = 1.10E-05), and the other three top entries were single organism process (GO:0044699; FDR = 3.10E-06), proteasome assembly (GO:0043248, FDR = 1.10E-05), and generation of precursor metabolites and energy (GO:0006091; FDR = 2.60E-05) (Figure 4B, Supplementary Table S3). The CC-related genes were enriched into 18 main GO terms, with cytoplasm, cell membrane, and membrane-bounded organelle representing the dominant CC categories (Supplementary Table S3). Using the KEGG database, the pathway enrichment analysis revealed that uniquely nonadditively expressed DEGs in Zheng58 × lx9801^*hlEW2b*^ vs. Zheng 58 × lx9801 were mainly enriched in carbon metabolism, proteasome, protein export, and biosynthesis of amino acids (Figure 5). Thus, the uniquely nonadditively expressed DEGs in Zheng 58 × lx9801^*hlEW2b*^ vs. Zheng 58 × lx9801 that are involved in these GO/KEGG categories (extensive biosynthetic and metabolic activities) may play important roles in maize ear development and early heterosis formation.

**Figure. 4.**
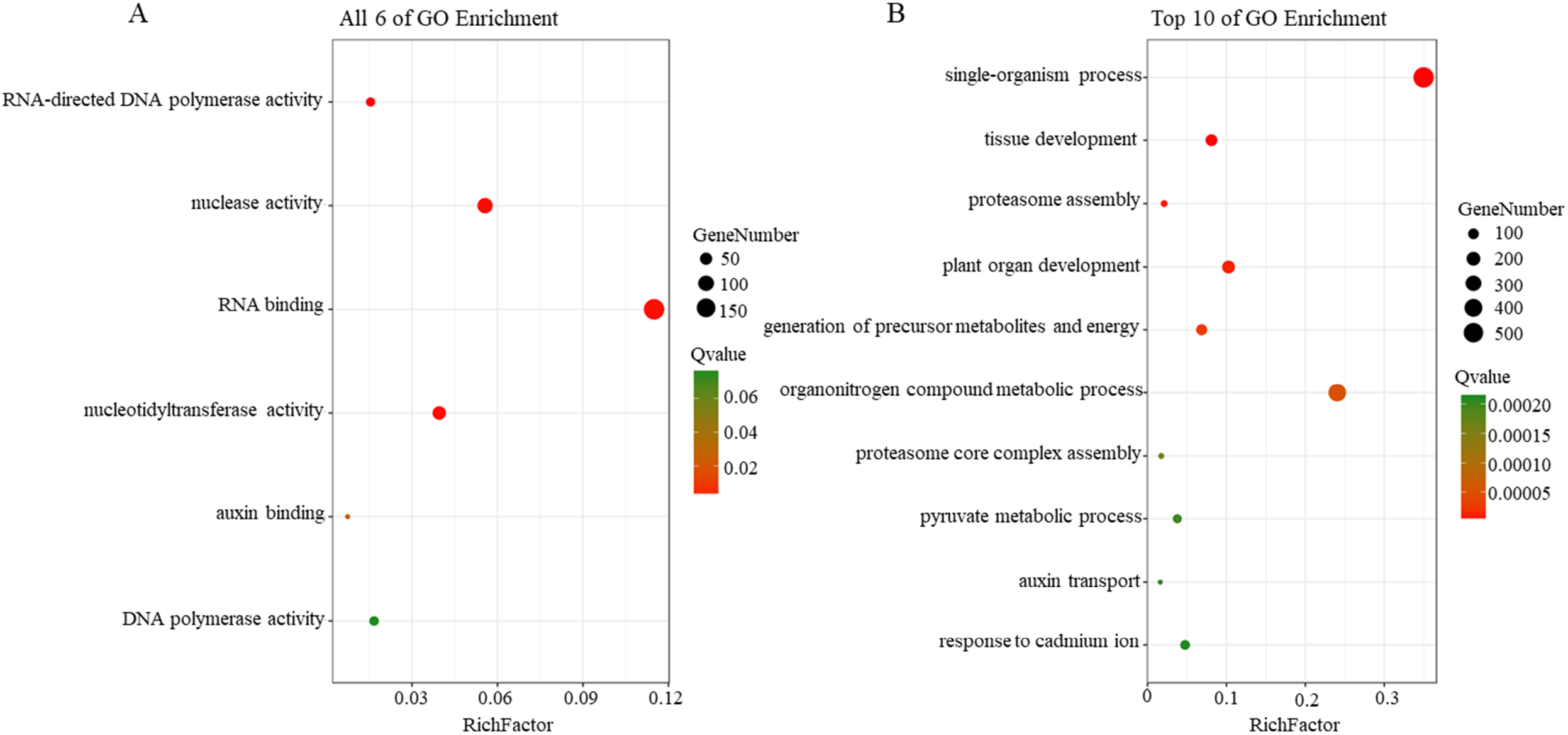
GO enrichment analysis of non-additive effects genes. A: Molecular functions GO enrichment result of non-additive expression genes selected from each hybrid. B: Biological process GO enrichment result of non-additive expression genes selected from each hybrid. Y-axis list the enrichment GO terms, X-axis represents rich factor. The smaller Q value, the higher significant level and more close to the red point. The more genes, the larger point.

**Figure. 5.**
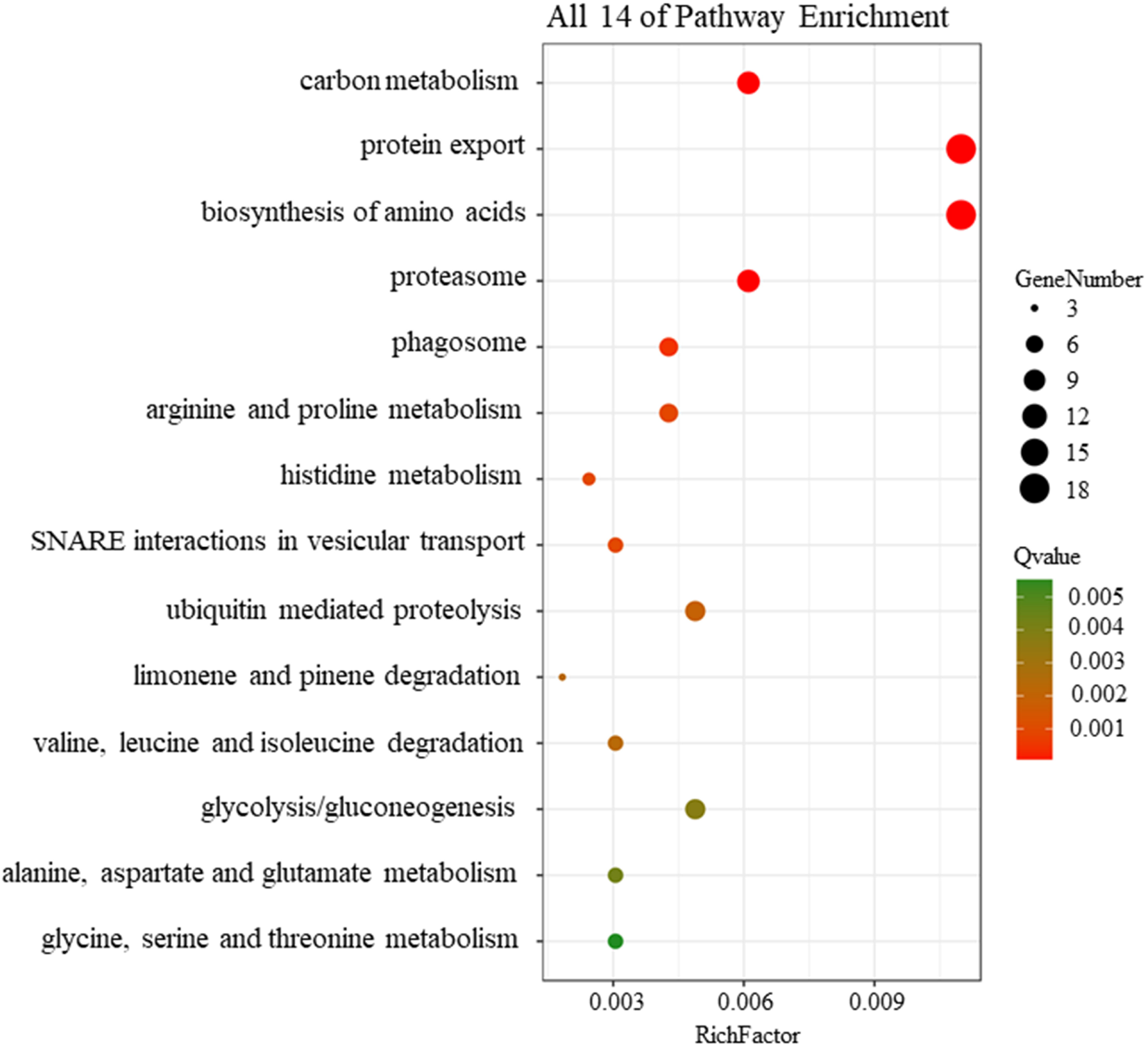
KEGG enrichment analysis of non-additive gene. Y-axis list the enrichment KEGG pathway, X-axis represents rich factor. The smaller Q value, the higher significant level and more close to the red point. The more genes, the larger point.

### Allele-specific expression analysis

Based on the single nucleotide polymorphism (SNP) information and specific allele sequencing data for the parents of hybrids, global allele-specific expression genes were identified in hybrids. A total of 362,410 and 360,791 polymorphic SNPs were identified between lx9801 and Zheng 58 and between lx9801^*hlEW2b*^ and Zheng58, respectively (Table 2). The specific expression levels (P) were defined as follows: the expression of the allele divided by the sum of the expression levels of the alleles representing the two parental genotypes. Thus, when P = 0, only the allele of lx9801 (Zheng58 × lx9801) or lx9801^*hlEW2b*^ (Zheng58 × lx9801^*hlEW2b*^) was expressed in the hybrid, and when P = 1, only the allele of Zheng58 was expressed. When P = 0.5, no allele-specific expression occurred. Furthermore, it was necessary to combine the SNP and hybrid allele read data to identify the ASE genes using the following criteria: the allele-specific expression level (P) is greater than 0.8 or less than 0.4; 2) the posterior probability is greater than 0.8; and 3) there are at least two SNPs on a single gene. In accordance with these standards, in the hybrids Zheng58 × lx9801 and Zheng58 × lx9801^*hlEW2b*^, 2263 genes (12.89% of 17,563 analyzed genes) and 2352 genes (14.65% of 16,059 analyzed genes) were identified as having significant allelic bias, respectively (Figure 6, Table 2). After comparison, 1310 (39.6%) ASE genes were identified in both hybrids, 1042 (31.5%) ASE genes were only expressed in Zheng58 × lx9801^*hlEW2b*^, and 935 (28.9%) ASE genes were only expressed in Zheng 58 × lx9801 (Supplementary Figure S1). The results indicated that different ASE genes expressed in different hybrids might greatly influence heterosis.

**Table 2.** Allele-specific gene expression genes in two hybrids.

| Sample | Protein coding genes | Significant ASEs | ASE genes ratio (%) |
| --- | --- | --- | --- |
| **Zheng58×lx9801** | 17563 | 2263 | 12.89 |
| **Zheng58×lx9801<sup>hIEW2b</sup>** | 16059 | 2352 | 14.65 |

**Figure. 6.**
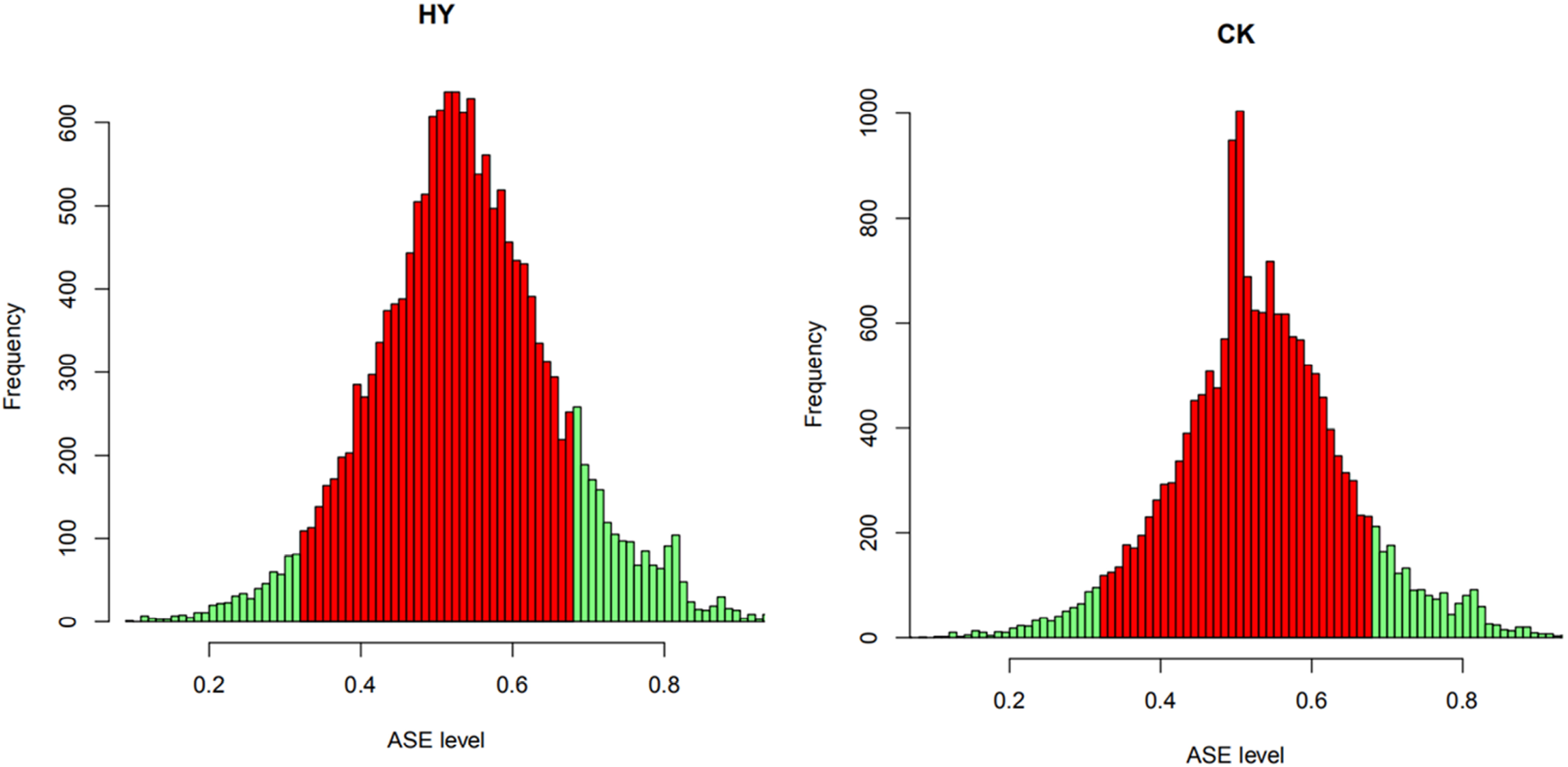
The distribution of ASE genes in hybrids. Green and red represent allele-specific expression genes and no allele-specific expression genes at the genome-wide level in Zheng58 × lx9801^*hlEW2b*^ (HY) and Zheng58 × lx9801 (CK).

In the present study, most of the ASE genes shared by Zheng58 × lx9801^*hlEW2b*^ and Zheng 58 × lx9801 were enriched in some basic biological processes, such as cellular macromolecular complex assembly, cellular component assembly, and organonitrogen compound metabolic processes. The unique ASE genes in Zheng 58 × lx9801^*hlEW2b*^ were most significantly enriched in single organism process, which was consistent with the GO analysis results of nonadditively expressed genes (Supplementary Table S4). Some classical genes, such as *WUS*, AINTEGUMENATA (*ANT*), and *KNOX*, which directly affect the fundamental formation of maize ear architecture, have been shown to be enriched in this GO term. Additionally, the specific ASE genes of Zheng58 × lx9801^*hlEW2b*^ were also significantly enriched in biological processes related to biosynthesis and catabolism (Supplementary Table S4), which implied that material metabolism provided support for a strong heterotic performance.

### Auxin involved in heterosis formation during maize ear development

In this study, the critical genes in the auxin response and synthetic transport pathways were screened based on the transcriptome data, and the transcription factors auxin response factor (*ARF*) and *APETALA*_*2*_, the auxin response genes Small Auxin Up Regulated (*SAURs*) and Gretchen Hagen 3 (*GH3*), as well as the auxin polar transport gene *brachyric2*, were all enriched in single organism process. Therefore, the expression levels of 38 *ARF* genes in the maize genome were analyzed in this study (Supplementary Figure S2A). *ARF1* and *ARF27* showed differential expression (both genes downregulated in Zheng58 × lx9801^*hlEW2b*^) and were also expressed nonadditively in two hybrids (F_1_ vs. MPV, FDR<0.05) (Figure 7, Table 3). In addition, the expression of the *brachyric2* gene was significantly higher in Zheng58 × lx9801^*hlEW2b*^ than in Zheng58 × lx9801 (Figure 7, Table 3), and it showed nonadditive expression patterns in Zheng58 × lx9801^*hlEW2b*^ and Zheng58 × lx9801 (Figure 7, Table 3). Notably, in the auxin tryptophan pathway, the flavin monooxygenases, *YUCCA*s, are critical enzymes that catalyze the rate-limiting step in auxin biosynthesis. Although *YUCCA* was not significantly enriched, it played a vital role in the auxin synthesis pathway. Of the nine maize *YUCCA* genes, only *YUCCA8* was significantly upregulated in Zheng58 × lx9801^*hlEW2b*^ compared with Zheng58 × lx9801 (Figure 7, Table 3, Supplementary Figure S2B), and it was expressed significantly higher and lower than the MPVs in hybrids Zheng58 × lx9801^*hlEW2b*^ and Zheng58 × lx9801, respectively (Table 3). This suggests that *YUCCA8* is nonadditively expressed in the two hybrids and shows positive heterosis in Zheng58 × lx9801^*hlEW2b*^. Thus, the differential expression of auxin-regulated genes might lead to changes in auxin levels and affect immature ear development in hybrids, resulting in heterotic performance difference between Zheng58 × lx9801^*hlEW2b*^ and Zheng58 × lx9801.

**Figure. 7.**
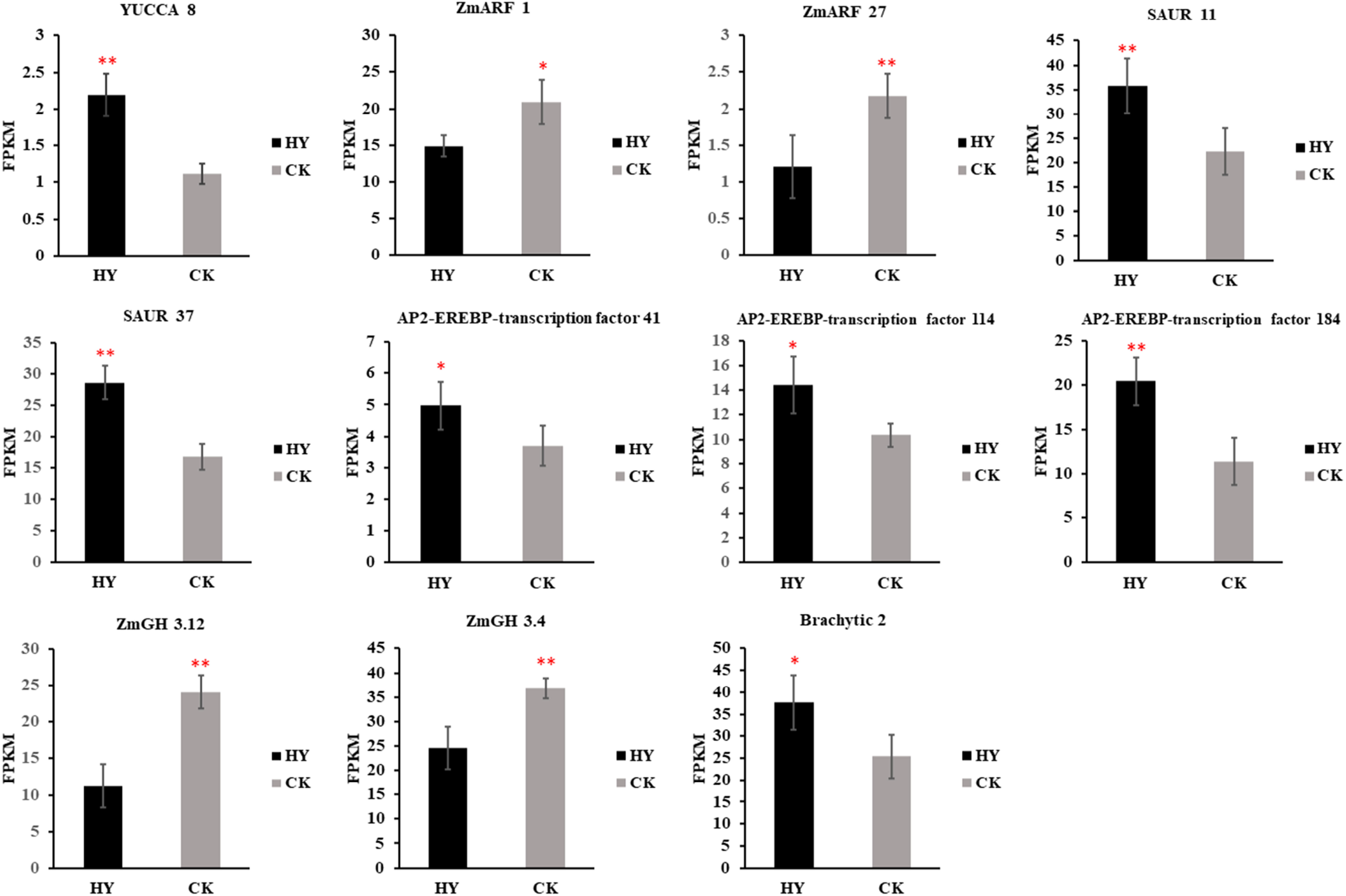
Expression analysis of auxin associated genes in the hybrids. Expression levels are presented as the FPKM average of three replicates ± standard error. HY: Zheng58 × lx9801^*hlEW2b*^; CK: Zheng58 × lx9801. * and ** indicate significant differences at FDR < 0.05 and FDR < 0.01, respectively.

**Table 3.** Heterosis effect analysis of auxin-related genes.

| GENE ID | Fold change from MPV, FDR ≤0.05 |  |  |  | GENE |
| --- | --- | --- | --- | --- | --- |
|  | HY | CK | lx9801<sup>hIEW2b</sup> | lx9801 | DESCRIPTION |
| Zm00001d011764 | 0.96 | -0.84 | -0.15 | -0.12 | *YUCCA 8* |
| Zm00001d051302 | 0.21 | 0.82 | -0.05 | 0.44 | *SAUR 11* |
| Zm00001d020605 | 0.48 | 0.80 | -0.50 | -2.1 | *SAUR 37* |
| Zm00001d031871 | 2.10 | 3.41 | 1.06 | 0.87 | *Brachytic 2* |
| Zm00001d007840 | 0.50 | 0.71 | -0.47 | -0.47 | *ANT 41* |
| Zm00001d018731 | 0.11 | 0.32 | -0.45 | -0.45 | *ANT 114* |
| Zm00001d034204 | 0.03 | 0.54 | -0.08 | -0.08 | *ANT 184* |
| Zm00001d031522 | -0.87 | -0.19 | 0.47 | 0.30 | *ZmARF 1* |
| Zm00001d045026 | -1.27 | -0.48 | -0.11 | -0.26 | *ZmARF 27* |
| Zm00001d006753 | -0.36 | 1.07 | 0.24 | -0.17 | *ZmGH 3.4* |
| Zm00001d022017 | -0.08 | 0.71 | 0.61 | 0.49 | *ZmGH 3.1* |

Note: 1) HY and CK represents Zheng58 × lx9801<sup>hIEW2b</sup> and Zheng58 × lx9801, respectively.
2) The genes involved in the table are significantly different from the MPVs of the respective hybrid combinations. Significant level threshold is FDR<0.05.
3) Red and green color indicate the up and down regulated fold change (FC) from the each of the two hybrid combinations MPVs.

## Discussion

### Overdominance effects may be the greatest contributors to heterosis

In recent years, many studies have shown that overdominance effects provide certain contributions to heterosis, and some genes producing overdominance effects have been identified. Krieger et al. (2010) found that the *SFT* gene was heterozygous for a functional allele and that a loss-of-function allele resulted in an overdominant performance for tomato yield. Additionally, heterotic loci with overdominant effects at the single-site level have been identified using genetic mapping populations (Hua et al., 2003). Unfortunately, these so-called overdominant genes or heterotic loci were only analyzed at a single gene or locus, without controlling for the complex genetic backgrounds of the materials themselves. Therefore, pseudo-overdominance and epistatic heterotic effects may exist at the genomic level. In this study, single-fragment substitution lines were used to eliminate the interference of genetic background. To analyze the heterotic effects of *hlEW2b*, the phenotypes of ear width in the test hybrid (Zheng58 × lx9801^*hlEW2b*^) and its parental inbred lines (Zheng58 and lx9801^*hlEW2b*^) were identified under multiple environmental conditions, and the heterotic loci *hlEW2b* had an overdominance effect on the heterosis of ear width.

### Nonadditive expression patterns play critical roles in the heterosis of ear width in maize

By analyzing the transcriptome data of hybrids and parents, the gene expression patterns can be divided into additive and nonadditive expression (Nadine et al., 2008). The increased activities of nonadditively expressed genes in Arabidopsis are related to the photosynthetic capacity promotion, as well as cell size and number, which indicates that nonadditively expressed genes may play important roles in the biomass heterosis (Fujimoto et al., 2012). Meyer et al. (2012) proposed that the nonadditively expressed genes can effectively improve the resource use of Arabidopsis seedlings and promote the enhancement of metabolic activities, which show strong heterosis in hybrids. However, in some cases, additively expressed genes are the greatest contributors to heterosis, or the combined expression of additive genes and nonadditive genes has the same effects on heterosis (Swanson-Wagner et al., 2006). The above differences may be caused by different experimental designs and statistical analyses. Here, a special experimental design was used to explore the relationship between gene expression pattern alteration and heterosis. The DEGs were identified from Zheng58 × lx9801^*hlEW2b*^ vs. Zheng 58 × lx9801, and the special DEGs in hybrid Zheng58 × lx9801^*hlEW2b*^ might have greatly influenced the performance of heterosis. Additionally, these nonadditively expressed genes from Zheng58 × lx9801^*hlEW2b*^ and Zheng58 × lx9801 were combined into one set, in which each gene showed a nonadditive expression pattern in at least one hybrid combination and displayed different heterotic performances between Zheng58 × lx9801^*hlEW2b*^ and Zheng58 × lx9801.

The GO enrichment analysis revealed that these nonadditively expressed genes were mainly overrepresented in basic biological processes related to growth and development (Figure 4A). According to functional genomics research, some key genes that regulate the transport and synthesis of auxin and affect the development of maize reproductive organs are significantly enriched in the processes of plant organ development (GO.0099402) and tissue development (GO.0009888). In the most highly represented GO categories (GO.0044699: single organism process), *MADS-transcription factor 1* and *9* genes, as well as *AGAMOUS* genes, were significantly enriched. The *MADS* genes are structurally conserved in monocot (such as maize) plants and dicot (such as Arabidopsis) plants, and mainly affect the formation of reproductive organs and regulates the conversion process between vegetative and reproductive growth (Ng and Yanofsky, 2001). The *AGAMOUS* genes are necessary for meristem development and play important roles in maintaining the direction of meristem development (Schmidt et al., 1993). Several transcription factor families have been significantly enriched, such as *NAC, MYB*, and *ARF*, which mainly function to influence the development of meristems and the formation of reproductive organs through the transcriptional regulation of plant growth and development (Hai et al., 2012; Ko et al., 2010). Among them, *ARF* is not only enriched in the single organism process, but also in the process of plant organ development. In plants, *AUXIN/INDOLE-3-ACETIC ACID* (*Aux/IAA) transcription factor* and *ARF* form a complex that is involved in auxin synthesis and ultimately affects maize ear development (Liscum and Reed, 2002). Additionally, in the processes of organ development (GO.0099402) and tissue development (GO.0009888), some important genes, such as *thick tassel dwarf1, fasciated ear2*, and *brachytic2*, that affect meristem differentiation were also significantly enriched. The expression of maize *fasciated ear2* promotes the differentiation of apical meristem, and its weak mutation leads to increases in IM size and kernel row numbers (Peter et al., 2013). The *thick tassel dwarf1* gene directly affects the development of IM, leading to the excessive proliferation of IM and causing malformations in female ear development (Peter et al., 2005). The *brachytic2* gene plays an important role in the polar transport of auxin (Wei et al., 2018). Thus, hormone regulation may also be a reason for the differential performance of heterosis in early stages of maize ear development. Additionally, nonadditively expressed genes are involved in a wide range of basic biosynthetic processes (map1230: Biosynthesis of amino acids) and material metabolism (map1200: Carbon metabolism; map3060: Protein export) (Figure 5), which are fundamental for producing the basic components and energy sources for heterosis formation in early maize ear development.

### ASE genes involved in maize ear development

The specific expression of alleles or the unbalanced expression of the two parental alleles in hybrids are important causes of heterosis (Guo et al., 2004; Shao et al., 2019). A competitive transcriptional relationship between two alleles may be the cause of the differential expression of genes in hybrids. The expressions of alleles can be divided into three modes: 1) two alleles simultaneously expressed at the same abundance levels in hybrids; 2) two alleles having competing expression at different abundance levels in hybrids; 3) one allele predominantly expressed and the other allele silenced in hybrids. From the perspective of heterosis-related studies, these patterns support the classical genetic hypotheses of dominance and overdominance (Shao et al., 2019). In this study, the proportion of ASE gene was almost equal between the two hybrids, which may result from them having the same female parent (Zheng58) and the male parents beings from near isogenic lines (lx9801 and lx9801^*hlEW2b*^). Although the number of ASE genes produced in each hybrid was limited, there was a slightly difference in the ASE genes between the two hybrids. The ASE genes shared by the two hybrids were enriched in some basic biological processes, such as cellular macromolecular complex assembly, cellular component assembly, and organonitrogen compound metabolic process. The enrichment of these basic biological processes may contribute to the performance of heterosis. Notably, consistent with the GO enrichment results for nonadditive genes, a GO enrichment with the special ASE genes in Zheng 58 × lx9801^*hlEW2b*^ revealed that the most significant item was also the single organism process (GO.0044699) (Figure 4B, Supplementary Table S4). Some critical genes (*WUS, ANT* and *KNOX*, etc.) related in inflorescence development were enriched in this term, which could involve in important processes of ear development. For instance, the *ANT* gene affects the formation of floral organs by regulating cell division and cell number (Mizukami and Fischer, 2000). The *WUS* gene plays an important role in determining the direction of cell differentiation and in regulating plant organ development (Farquharson, 2015), and the *KNOX3* and *5* genes are involved in the maintenance of meristem activity and the formation of the initial floral organ developmental pattern (Hake et al., 2004). These special ASE genes in Zheng58 × lx9801^*hlEW2b*^ may be important reasons for it having stronger heterosis than Zheng58 × lx9801.

### Auxin may contribute to the hybrid heterosis of ear development in maize

Morphological analyses have shown that the most obvious feature of the transformation from vegetative to reproductive growth is the rapid increase in the size of the IM. The cells located in the central zone of the IM have typical stem cell characteristics, and they can initiate and determine the developmental processes of maize ears (Vollbrecht and Schmidt, 2009). The *CLV–WUS* negative feedback loop may affect the development of IM by regulating the relationship between the proliferation of stem cells and the differentiation activities of tissues and organs. In this pathway, the *WUS* gene, located in the organizing center of the meristem, activates the expression of the signal molecule *CLV3*, which is sensed by *CLV1–CLV2* and they form a complex. This complex positively regulates the expression of the *WUS* gene and promotes stem cell proliferation. However, *CLV1, CLV2*, and *CLV3* in the noncomplexed state inhibit *WUS* gene expression and form a dynamically balanced negative feedback loop that affects the IM differentiation processes (Somssich et al., 2016). The genes involved in the *CLV–WUS* negative feedback loop have also been cloned in maize, including thick *tassel dwarf1* and *fasciated ear2*, which are homologs of *CLV1* and *CLV2*, respectively. The homologous gene of signal molecule *CLV3* in maize is *COMPACT PLANT2* (Bommert et al., 2013; Peter et al., 2005; Taguchi-Shiobara et al., 2001). Moreover, the expression of the *WUS* gene may be induced by auxin, and the abundance of *WUS* gene expression is positively correlated with the auxin gradient, which implies that auxin regulation is critical for IM development (Su et al., 2009).

In the present study, several critical genes involved in inflorescence development and auxin metabolism have been identified. Moreover, these genes were differentially expressed between the two test hybrids, Zheng58 × lx9801^*hlEW2b*^ and Zheng58 × lx9801, and showed nonadditive expression or ASE patterns in hybrids, which may lead to the differential performance of heterosis between the hybrids. In the IM, the upregulated *CLV1* [*tassel dwarf1*, Zheng 58 × lx9801^*hlEW2b*^ (HY) vs. Zheng 58 × lx9801 (CK), FDR = 1.66E-02, log_2_ (HY/CK) = 0.57] and *CLV2* [*fasciated ear2*, HY vs. CK, FDR = 2.20E-02, log_2_ (HY/CK) = 0.54] combine with the *CLV3* [*COMPACT PLANT2*, HY vs. CK, FDR = 5.14E-01, log_2_ (HY/CK) = 0.29] to form a complex, which enhances the *WUS* [HY vs. CK, FDR = 1.07E-03, log_2_ (HY/CK) = 1.75] gene expression activity and maintains the size of the stem cell population to positively regulate IM development (Figure 8). In addition, Rodriguez et al. (2016) and Perales et al. (2016) proposed a model in which *WUS* dimers negatively regulate *CLV3* expression in the organizing center, and *WUS* monomers positively regulate *CLV3* expression in the central zone. This may explain why *CLV3* does not show a significant upregulation in the dominant hybrid of Zheng58 × lx9801^*hlEW2b*^.

**Figure. 8.**
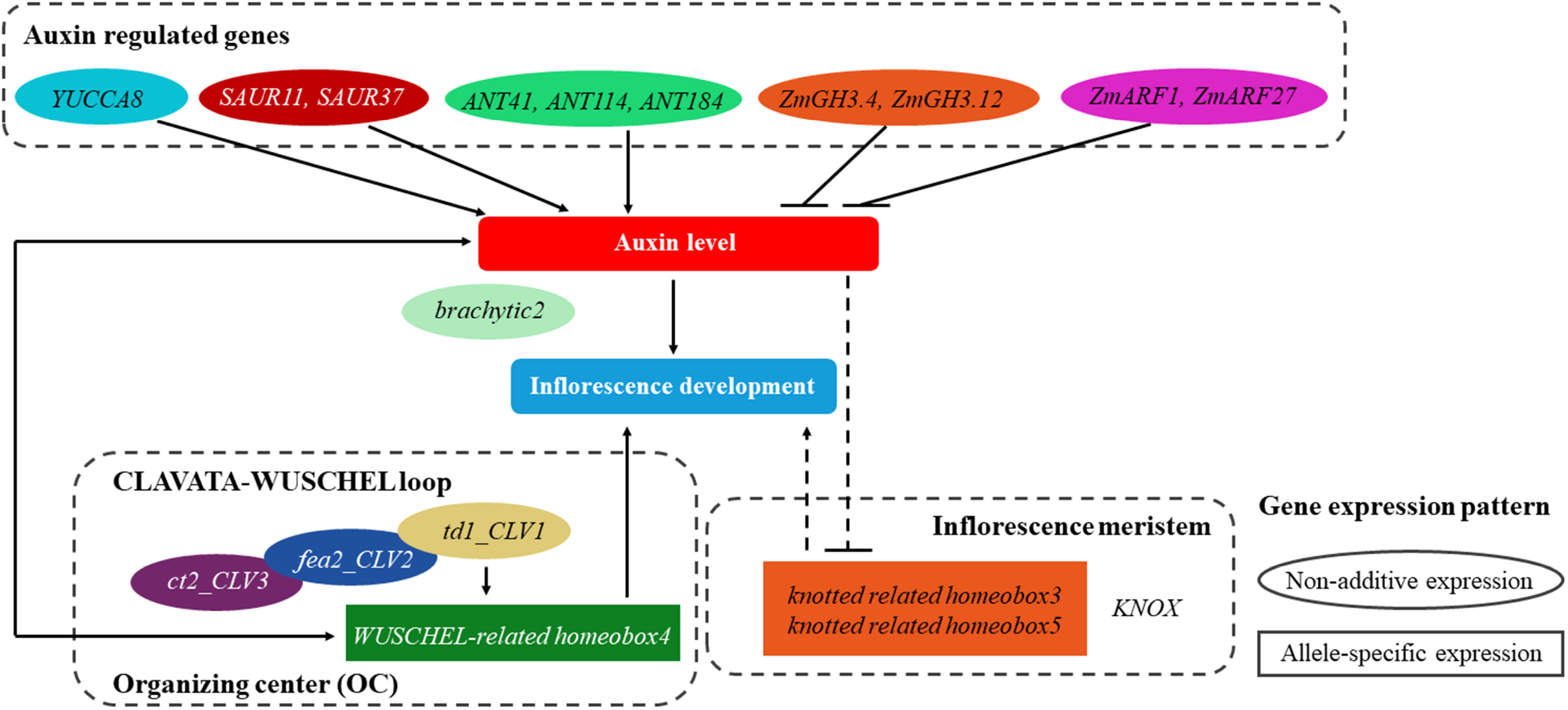
Predicting the regulatory process of heterosis formation during inflorescence meristem period. Auxin may contribute to hybrid heterosis in maize ear development. Non-additive expression and ASE expression may fine-tune the expression levels of crucial genes that control auxin metabolism and IM development to an optimal status, which may be responsible for the maize ear heterosis formation in hybrids. Oval represents non-additive expression genes. Rectangles represents allele-specific expression (ASE) genes.

Local auxin gradients may play roles in the establishment of organs in the IM (Benková et al., 2003). The tryptophan pathway is critical for auxin synthesis. In this pathway, *YUCCA* is the rate-limiting enzyme that catalyzes auxin biosynthesis (Youfa et al., 2006). Of the nine *YUCCA* genes in maize, *YUCCA8* was upregulated in the hybrid Zheng58 × lx9801^*hlEW2b*^, compared with in the control hybrid Zheng58 × lx9801, which implies that the auxin accumulation level in Zheng 58 × lx9801^*hlEW2b*^ may be higher than in Zheng58 × lx9801. We also analyzed the auxin-responsive genes, *SAUR11, SAUR37*, and *ANT*, that affect cell proliferation and floral organ development, and these genes were all upregulated in Zheng 58 × lx9801^*hlEW2b*^. The *SAUR* genes are regarded as early and primary auxin-responsive genes involved in the auxin signal transduction pathway. The overexpression of *SAUR* genes may significantly increase the cell proliferation rates in plants (Ren and Gray, 2015). The *ANT* gene is a transcription factor containing an *APETALA*_*2*_ domain, and it can affect the development of floral organs by regulating the auxin level (Krizek, 2009). The upregulated expression of these auxin-regulating genes may be responsible for the increase of IM width in the hybrid Zheng 58 × lx9801^*hlEW2b*^. The *brachyric2* gene is an auxin export carrier that participates in the polar transport of auxin and affects auxin transport (Knoeller et al., 2010). The upregulated expression of this gene in Zheng 58 × lx9801^*hlEW2b*^ (Figure 8) is conducive to the local accumulation of auxin in meristems, thereby ensuring that the meristem maintains a strong meristematic capability. Additionally, IAA-amide synthase, encoded by the auxin negative regulator *GH3*, converts active auxin into an inactive state (Ostrowski and Jakubowska, 2013). The downregulation of the *GH3* gene results in a large accumulation of active auxin in Zheng58 × lx9801^*hlEW2b*^. At low auxin concentrations, *ARF* proteins form heterodimers with *Aux/IAA* proteins, which repress the transcriptional activities of *ARF*s (Wang and Estelle, 2014). Therefore, it is inferred that the decrease in the *ARF* transcriptional activity of Zheng58 × lx9801^*hlEW2b*^ leads to the accumulation of auxin in the IM. Furthermore, high relative auxin concentrations in the primordial region appear to facilitate organ initiation through the downregulation of *KNOX* expression (Heisler et al., 2005). In this study, the alteration of the expression levels of several critical auxin metabolism genes may lead to high auxin concentrations in the IM, which possibly changes the *KNOX* gene expression pattern and promotes primordium initiation (Figure 8).

In present work, the transcriptome analysis determined that the expression of genes involved in auxin metabolism in Zheng58 × lx9801^*hlEW2b*^ was better coordinated than in the control hybrid Zheng58 × lx9801, and this was more beneficial to the synthesis and accumulation of auxin. Additionally, the gene expression pattern analysis showed that nonadditive expression and ASE might be the causes of differential gene expression between hybrids. It implied that nonadditive expression and ASE may fine-tune the expression levels of crucial genes that control auxin metabolism and IM development to an optimal stats, and this may be responsible for the heterosis of maize ear formation in hybrids.

## Acknowledgements

National Natural Science Foundation of China (31971961, 31871641), National Natural Science Foundation of Henan Province (202300410215), the Key Technologies R & D Program of Henan Province (212102110061, 212102110043), Basic and Frontier Technology Research Program of Henan Province (162300410165).

## Data availability

The data that support the findings of this study are openly available in “NCBI” at https://submit.ncbi.nlm.nih.gov/

## Author contributions

J.T. and W.L. designed the experiments and together with, X.S, X.S. and Z.G. carried out data analysis, M.W., X.G., L.Y., X.Q. and H.X. carried out maize morphological investigation and collected samples, Y.X. visualization of charts. X.S. and J.T. was mainly involved in preparation of manuscript. All authors reviewed and approved the final manuscript.

